# *In-silico* analysis of Myocardial Infarction-related missense SNPs to identify novel biomarkers to predict susceptibility

**DOI:** 10.1101/2023.11.07.565946

**Authors:** Fiza Faris Tarlochan, Saad Rasool

## Abstract

Myocardial Infarction (MI), commonly known as a heart attack, stands as a formidable global health challenge, responsible for a substantial burden of morbidity and mortality. This study embarked on a comprehensive exploration of the genetic underpinnings of MI, recognizing the pivotal role of genetic factors in determining an individual’s susceptibility to this life-threatening condition. The objective of our research was to investigate missense single nucleotide polymorphisms (SNP) associated with MI to determine whether the changes in amino acid sequences have potential implications for the risk of MI. Employing a multifaceted approach, we leveraged an array of computational tools and databases to scrutinize specific missense SNP and meticulously analyzed their potential effects on protein structure stability and function. Our analysis has confirmed a total of 4 missense SNP in ALDH2, APOE, IGFBP1, and PCSK1 genes to be damaging to protein structure and hence, the function. An extensive literature review was then performed to determine the functional roles of these genes in the regulation of the cardiac system-related pathways. Our analysis confirmed that all 2 of these genes are directly involved in pathways related to the cardiac system, while the other 2 genes play other roles. We have further analyzed their interactions and underlying biological processes to determine their potential role in the incidence of MI. These findings collectively offer a profound understanding of the intricate genetic landscape underlying MI. They not only enhance our comprehension of the multifaceted genetic factors influencing MI susceptibility but also set the stage for future experimental investigations. Importantly, these insights hold the potential to guide future research and the development of therapeutic strategies, to improve the prevention and management of this critical cardiovascular condition.

## 1. Introduction

Myocardial infarction (MI), commonly known as a heart attack, is a devastating cardiovascular event characterized by the occlusion of coronary arteries, resulting in insufficient blood supply to the heart muscle (Saleh M., Ambrose JA., 2018). It represents a major global health concern, accounting for a significant proportion of morbidity and mortality worldwide (Kim SJ., 2021). MI poses a substantial burden on individuals, families, and healthcare systems, necessitating extensive research to unravel its complex etiology, identify potential therapeutic targets, and develop effective strategies for management and prevention.

MI arises from the interplay of various risk factors, including age, gender, hypertension, dyslipidemia, smoking, diabetes, and a family history of cardiovascular disease (Saleh M., Ambrose JA., 2018). In recent years, it has become increasingly evident that genetic factors play a crucial role in modulating an individual’s susceptibility to MI (Dai X., et al., 2016). Therefore, understanding the molecular mechanisms underlying MI is of paramount importance as it will allow researchers and clinicians to gain critical insights into its pathophysiology, identify novel therapeutic targets, and develop interventions aimed at reducing the occurrence and severity of MI.

One promising avenue of research in understanding the genetic underpinnings of MI involves the investigation of Single Nucleotide Polymorphisms (SNPs), which are the most common form of genetic variation, representing single base pair changes in the DNA sequence occurring throughout the human genome. These genetic variations can impact gene expression, protein function, and biological pathways, thereby influencing an individual’s susceptibility to various diseases, including cardiovascular disorders such as MI. For our research, we are interested in missense SNPs as they involve changes in amino acid sequences. We hypothesize that changes in sequences can lead to changes in protein structure and function.

Association between SNPs and the susceptibility to MI. Notably have been explored in the past. Investigations in the Han Chinese population have revealed significant association between genetic variations, such as rs1764391 and rs1122608, and the risk of MI (Li J., et al., 2020, Chen QF., et al., 2018). In addition, certain genotypes of rs1122608 have also demonstrated a notable correlation with the time of MI diagnosis (Chen QF., et al., 2018). Furthermore, the presence of SNPs in CLOCK (rs6811520, rs13124436) and ARNTL (rs3789327, rs12363415) has also pointed towards potential interactions between the circadian rhythm and susceptibility to in MI (Škrlec I., et al., 2020).

Moreover, investigations have revealed that carriers of GG genotype of rs3480 exhibited significantly elevated levels of Troponin I and triglycerides. Similarly, carriers of GA genotype of rs726344 have displayed significantly increased levels of CKMB, total cholesterol, LDLc, HDLc, Troponin I, and triglycerides when compared to individuals with other genotypes, which further increases the risk of MI (Badr EA., et al., 2020). Furthermore, rs1143634 C/T in IL1B is also associated with blood lipid levels, including low-density lipoprotein (LDL) and total cholesterol (TC), in the Eastern Chinese Han population, potentially influencing MI risk (Pan Q., et al., 2021). Conversely, in a population from Western Siberia (Russia), the presence of the G allele of rs708272 has been associated with lower levels of high-density lipoprotein cholesterol and a higher index of atherogenicity, implicating this SNP in the development of MI (Semaev S., et al., 2019).

In this study, we harnessed a range of bioinformatics tools to pinpoint missense SNPs linked to myocardial infarction. We characterized the amino acid alterations, evaluated their impact on protein stability, and explored their protein-protein interactions. This comprehensive analysis sheds light on the influence of these SNPs on the development of myocardial infarction.

## 2. Methods

### 2.1 Data Curation and SNP Identification

Relevant SNPs were found using the search term “Myocardial Infarction’’ in the NHGRI-EBI GWAS database (Buniello et al., 2019). The information in the generated file includes replication sample size, chromosomal area, mapped gene, SNPs rs code, and p value.

### 2.2 Gene-level functional analysis of mapped SNPs

g:Profiler was used to undertake functional analysis of missense SNPs linked to genes. This program evaluates the specified target genes using the WikiPathway, KEGG, Reactome, and Gene ontology databases simultaneously and displays the findings with significant p values (p < 0.05) (Uku et al., 2019).

### 2.3 Amino acid change prediction

Sorting Intolerant from Tolerant (SIFT) algorithm was employed to prioritize and classify the potential impact of Single Nucleotide Polymorphisms (SNPs). SIFT is a widely used computational tool designed to predict the functional consequences of amino acid substitutions in proteins. The SIFT algorithm utilizes sequence conservation information to assess the likelihood of an amino acid change being deleterious or tolerated. To sort the SIFTs, we input the SNP rs-identifier information obtained from the mapped SNPs into the SIFT tool. The output of SIFT provided a sift score ranging from 0 to 1, where scores below a predefined threshold were considered potentially damaging, while scores above the threshold were deemed tolerant. By leveraging the SIFT algorithm, we were able to prioritize the SNPs based on their predicted functional impact, aiding in the identification and selection of functionally relevant SNPs for further analysis in our study.

### 2.4 SNP Pathogenicity Prediction

Sequence-based analysis was conducted employing PredictSNP, which combines various prediction tools such as PhD-SNP, PolyPhen 1 and 2, SIFT, and SNAP (Choi and Chan, 2015; Bendi et al., 2014). This analysis aimed to assess the pathogenicity of missense single nucleotide polymorphisms (missense SNPs). The outcomes were classified as either benign or pathogenic/deleterious based on established databases.

### 2.5 Protein Domain Architecture

The InterPro platform was employed to predict the protein function by submitting the amino acid sequences of deleterious genes. InterPro employs predictive models sourced from various databases to conduct functional analysis, encompassing molecule families, domains, essential sites, and other features (McDowall & Hunter, 2011). The data regarding protein domains was then utilized to create diagrams illustrating domain architecture using Inkscape Vector graphics software (Yuan et al., 2016).

### 2.6 Assessing the protein stability of deleterious genes

The stability of four harmful genes’ structures was assessed using DynaMut, mCSM, DUET, and Missense3D. DynaMut investigates and visualizes protein dynamics by analyzing the impact of mutations on stability through changes in vibrational entropy (Rodrigues et al., 2018). The mCSM server predicts alterations in protein stability due to specific mutations and provides molecular visualization of the protein structure (Pires et al., 2014). Additionally, the DUET server was employed to observe changes in protein stability resulting from mutations. In contrast to other computational tools, DUET offers an integrated analysis of protein stability by combining results from both mCSM and SDM (Pires et al., 2014). Finally, Missense3D predicts structural modifications arising from amino acid mutations (Ittisoponpisan et al., 2019).

### 2.7 Examining molecular pathways alongside protein partners associated with deleterious genes

The String Database (Szklarczyk et al., 2023) was employed to identify both known and predicted interacting partners of all deleterious proteins. Additionally, we retrieved enriched information on biological processes and pathways to discern the downstream effects of each interaction.

## 3. Results

### 3.1 Identification of missense SNPs associated with Myocardial Infarction

A total of 416 SNPs associated with myocardial infarction were identified through the NHGRI-EBI GWAS catalog. These SNPs were classified based on their variant effects using gProfiler’s gSNPense and were then organized and analyzed in Microsoft Excel. Figure 2 provides the frequencies of these variant effects. SNPs linked to myocardial infarction were detected in diverse regions, including non-coding transcripts, non-coding transcript exons, introns, NMD transcripts, and the 3’ untranslated region (3’UTR). Specifically, 233 SNPs were located in intronic regions, 144 in non-coding transcript regions, 80 in NMD transcripts, 25 were identified as missense, and only 4 were synonymous SNPs. Given that the focus of this study is to examine amino acid changes resulting from missense SNPs, only the 25 missense SNPs were further evaluated in various computational databases.

**Figure 1.**
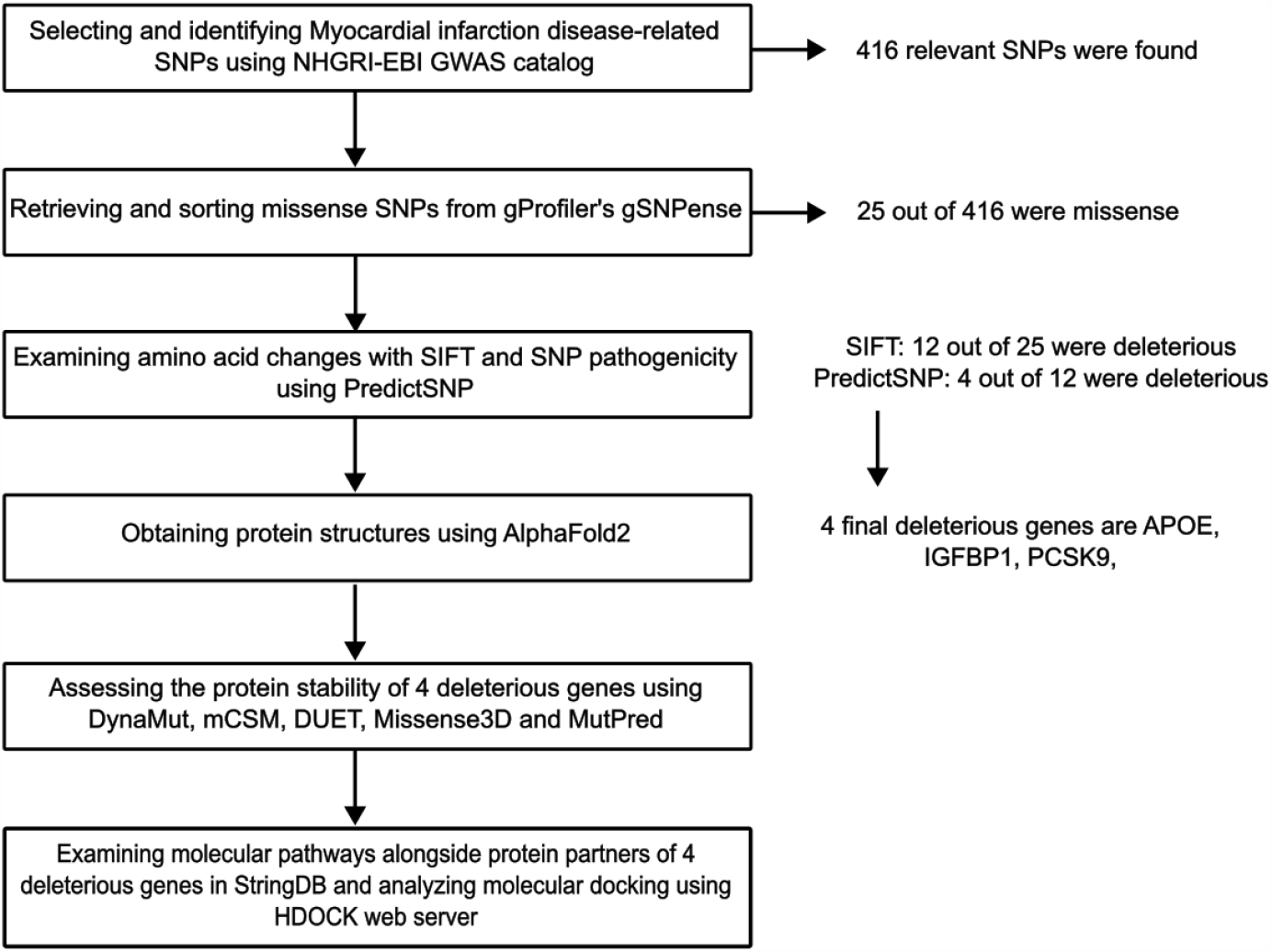
Project Workflow

**Figure 2.**
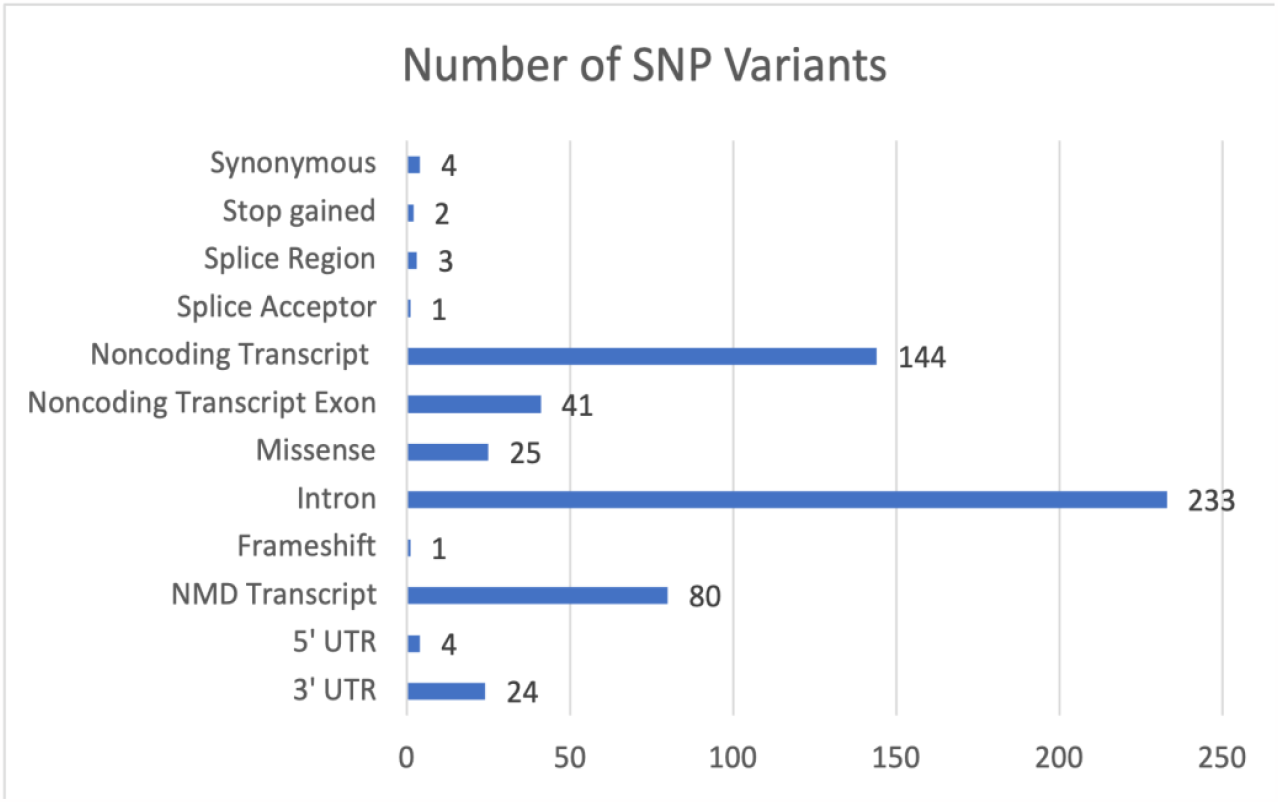
Frequency of each type of SNP variants associated with Myocardial Infarction

**Figure 3.**
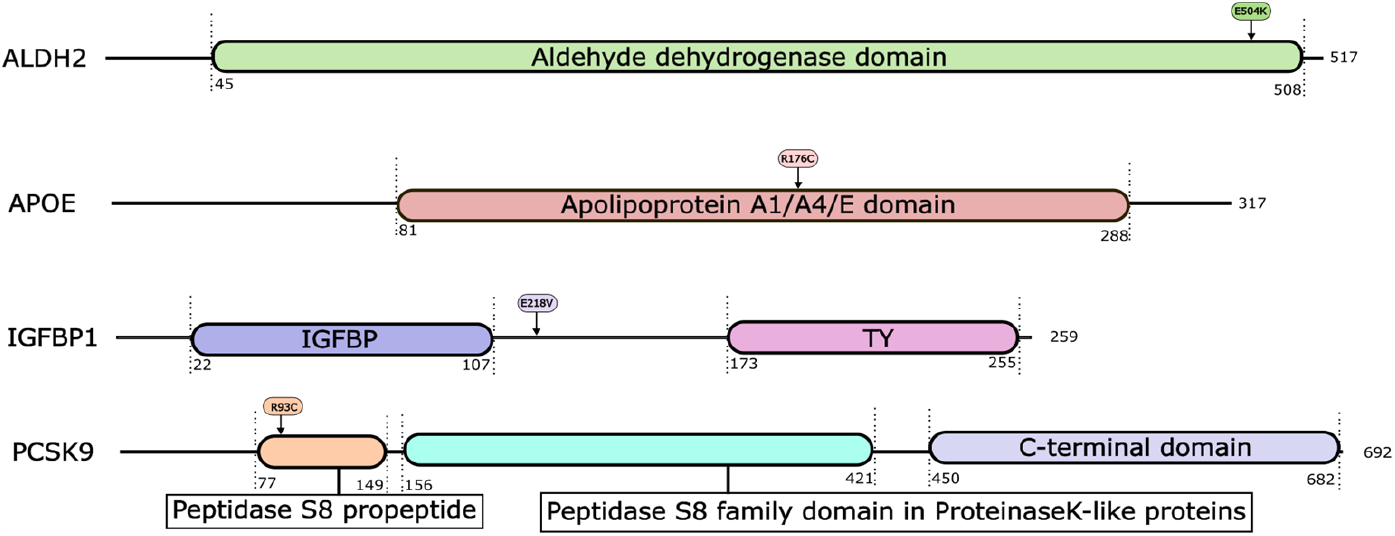
Domain architecture for deleterious genes with their respective amino acid substitution.

### 3.2 Mapping of missense SNPs and their respective amino acid changes

Out of the 25 missense SNPs, the SIFT prediction tool identified 12 missense SNPs as deleterious. The SIFT median scores for the identified deleterious genes ranged from 2.5 to 3.47. Additionally, SIFT provided the corresponding amino acid substitutions and the gene name.

### 3.3 Pathogenicity prediction of missense SNPs

The pathogenicity of all 12 missense SNPs, along with their corresponding amino acid substitutions, was predicted using PredictSNP, which provides prediction from a variety of programs, such as MAPP, PhD-SNP, etc. The selection of deleterious genes was made based on consensus predictions from all programs, and genes with unanimous deleterious predictions were earmarked for subsequent analysis. ALDH2, APOE, IGFBP1, and PCSK9 were singled out as deleterious genes to undergo further analysis with additional computational tools.

### 3.4 Analyzing the stability of protein structure

DynaMut, mCSM, SDM, and DUET servers provided ΔΔG values in kcal/mol for amino acid changes associated with ALDH2, APOE, IGFBP1, and PCSK9. Negative ΔΔG values indicate a decrease in stability, while positive ΔΔG values indicate an increase in stability. ALDH2 and APOE exhibited decreased protein stability due to the amino acid substitutions E504K and R176C, respectively. For PCSK9, the consensus from most servers predicted a decrease in protein stability, although one server indicated an increase. Regarding IGFBP1, half of the servers predicted decreased stability, while the other half predicted increased stability (Table 2).

**Table 1.** 12 missense SNPs related to Myocardial infarction disease, SNP identifier, amino acid substitutions, and the pathogenicity analysis from PredictSNP

| SNP ID | Gene Name | Amino Acid Change | MAPP | PhD-SNP | PolyPhen-1 | PolyPhen-2 | SIFT | SNAP |
| --- | --- | --- | --- | --- | --- | --- | --- | --- |
| rs199921354 | VPS33B | **R107Q** | 79% | 83% | 67% | 70% | 61% | 61% |
| rs115287176 | TMOD4 | **R277W** | 65% | 45% | 74% | 65% | 65% | 50% |
| rs146092501 | COL6A3 | **E1386K** | 66% | 68% | 67% | 63% | 67% | 61% |
| rs192210727 | LPHN3 | **R580I** | - | 68% | 67% | 47% | 79% | 58% |
| rs671 | ALDH2 | **E504K** | 64% | 68% | 59% | 41% | 79% | 56% |
| rs11601507 | TRIM5 | **V112F** | 73% | 58% | 67% | 64% | 45% | 58% |
| rs116843064 | ANGPTL4 | **E40K** | - | 55% | 59% | 68% | 53% | 55% |
| rs6761276 | IL1F10 | **I44T** | 63% | 66% | 59% | 63% | 43% | 55% |
| rs7412 | APOE | **R176C** | 86% | 83% | 74% | 81% | 53% | 62% |
| rs13306206 | APOB | **P955S** | 78% | 83% | 67% | 63% | 75% | 71% |
| rs117054298 | IGFBP1 | **E218V** | 57% | 86% | 67% | 47% | 79% | 56% |
| rs151193009 | PCSK9 | **R93C** | 74% | 61% | 74% | 68% | 53% | 62% |

Deleterious gene: Neutral gene:

**Table 2.**
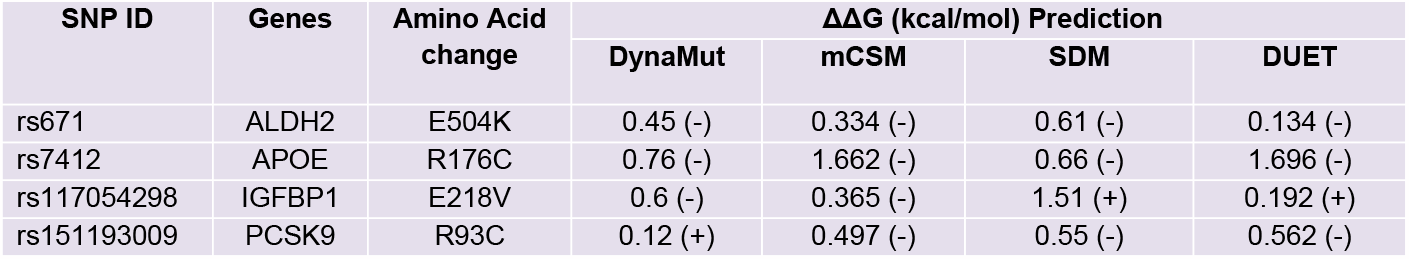
Protein structure stability results from four deleterious genes evaluated by DynaMut, mCSM, SDM, and Duet. The positive and negative signs refer to stabilizing and destabilizing respectively.

| SNP ID | Genes | Amino Acid change | $\Delta\Delta G$ (kcal/mol) Prediction | | | |
| --- | --- | --- | --- | --- | --- | --- |
|  |  |  | DynaMut | mCSM | SDM | DUET |
| rs671 | ALDH2 | E504K | 0.45 (-) | 0.334 (-) | 0.61 (-) | 0.134 (-) |
| rs7412 | APOE | R176C | 0.76 (-) | 1.662 (-) | 0.66 (-) | 1.696 (-) |
| rs117054298 | IGFBP1 | E218V | 0.6 (-) | 0.365 (-) | 1.51 (+) | 0.192 (+) |
| rs151193009 | PCSK9 | R93C | 0.12 (+) | 0.497 (-) | 0.55 (-) | 0.562 (-) |

### 3.5 Assessing the protein-protein interactions

Utilizing StringDB, we uncovered partner proteins associated with deleterious genes, delineating their molecular interactions, linked biological processes, and pathways. Our findings reveal that ALDH2 has three protein partners—ALDH6A1, ALDH1B1, and AGXT2—each engaging with the catalytic domain. Furthermore, APOE’s associations are with LDLR1 at its N-terminal region and LDLR2. IGFBP1’s interactions of IGF-1 at the N-terminal domain and both N and C-terminals for IGF-2. Lastly, PCSK9 engages with low-density lipoprotein receptors (LDLRs) and ANXA2.

## 4. Discussions

MI represents a critical cardiovascular condition with a substantial global impact. Investigating the genetic basis of MI, particularly through the study of SNPs, is essential for unraveling the complex mechanisms involved in its pathogenesis. The exploration of SNPs offers opportunities to identify at-risk individuals, improve risk stratification, and develop personalized treatment strategies. In this study we started with identifying all SNPs associated with Myocardial Infarction and shortlisted them to 4 genes including ALDH2, APOE, IGFBP1 and PCSK1, based on their impact on protein structure stability. We have further studied their interactions and underlying biological processes to determine their potential role in incidence of MI.

### ALDH2

ALDH2, also known as aldehyde dehydrogenase 2, is a pivotal enzyme with a vital role in aldehyde metabolism, particularly in the breakdown of acetaldehyde. A primary function of ALDH2 is the detoxification of acetaldehyde, a toxic byproduct generated during the breakdown of ethanol (alcohol) in the liver by alcohol dehydrogenase (Coudert E et al., 2023).

Beyond its well-established role in alcohol metabolism, ALDH2 has garnered attention for potential cardioprotective effects (Budas, G. R. et al, 2009). The enzyme is expressed in the cardiovascular system, and its activity has been linked to a reduced risk of cardiovascular diseases (Budas, G. R. et al, 2009). This protective effect is believed to stem from ALDH2’s involvement in aldehyde metabolism.

Our computational analysis revealed a missense single nucleotide polymorphism (missense SNP) in ALDH2, resulting in the amino acid change 5504K, which predicts a decrease in the protein’s stability. This compromised stability holds the potential to adversely impact the efficient metabolism of aldehydes, raising concerns about potential disruptions in the detoxification process and the cardioprotective property. Understanding the ramifications of this genetic variant on ALDH2 function provides valuable insights into the intricate interplay between enzyme stability and aldehyde metabolism.

Moreover, ALDH2 interacts with ALDH6A1, ALDH1B1, and AGXT2. The commonality among these genes lies in their coding for enzymes involved in various aspects of amino acid metabolism (Szklarczyk D,. et al, 2015). This interconnected network of enzymes underscores the significance of their collaborative roles in maintaining metabolic homeostasis and highlights the potential impact of genetic variations on these intricate pathways.

### APOE

Apolipoprotein E (apoE) is a pivotal player in lipid metabolism, crucial for the transport of cholesterol and other lipids in the bloodstream (Coudert E et al., 2023). Mutations in the APOE gene, responsible for coding apoE, can induce structural and functional alterations in the protein. Our computational analysis identified a missense single nucleotide polymorphism (missense SNP) in APOE, resulting in the amino acid change R176C, predicting a decrease in the protein’s stability. This reduced stability has the potential to impair the efficient transport of lipids by lipoproteins, leading to disruptions in lipid metabolism.

The diminished stability of apoE can potentially disrupt the effective transport of lipids by lipoproteins, resulting in disturbances to lipid metabolism. Of note is the critical interaction between apoE and the LDL receptor, encompassing both LDR1 and LDR2. This interaction is essential for the cellular uptake of lipoproteins (Szklarczyk D,. et al, 2015). A reduction in stability may adversely affect the binding affinity of apoE to these receptors, potentially leading to alterations in cellular recognition and the uptake of lipoproteins. The ultimate result is a potential reduction in the efficiency of lipoprotein clearance and lipid metabolism. These inefficiencies may significantly contribute to the development of conditions such as atherosclerosis, characterized by the accumulation of plaque within arteries, and other lipid-related disorders. Accumulation of lipids, a recognized factor in the pathogenesis of myocardial infarction, becomes especially critical when plaque rupture occurs.

Moreover, findings by Hopkins et al. underscore the significance of impaired removal of triglyceride-rich lipoproteins (TGRL), leading to the accumulation of abnormal TGRL remnants, known as β-VLDL. Elevated levels of these remnants contribute to lipid deposition in various conditions, including tuberous xanthomas, atherosclerosis, premature coronary artery disease, and early myocardial infarction (Hopkins PN,. et al., 2014). Investigating APOE in vivo and comprehending its role in lipid transport and metabolism can unveil valuable insights into the intricate mechanisms underlying these pathological processes.

### IGFBP1

Insulin-like growth factor-binding protein 1 (IGFBP-1) is a member of the insulin-like growth factor-binding protein family (Coudert E et al., 2023). This protein is primarily known for its role in regulating the activity of insulin-like growth factor 1 (IGF-1), a critical factor involved in cell growth, development, and metabolism (U.S. National Library of Medicine, 2018). IGFBP-1 is a protein that binds to IGF-1 with high affinity, forming a complex that modulates the bioavailability and activity of IGF-1 (Bae, J. H., et al., 2013). IGFBP-1 plays a crucial role in regulating the effects of IGF-1 by controlling its distribution and availability in the bloodstream. When IGFBP-1 binds to IGF-1, it can influence the transport of IGF-1 to target tissues and modulate its interaction with IGF-1 receptors.

According to the computational analysis, there is a 50% chance of the E218V missense SNP destabilizing the protein. Destabilization of IGFBP-1 may result in decreased binding affinity or increased dissociation of IGF-1 from IGFBP-1, leading to higher levels of free, biologically active IGF-1. Elevated levels of free IGF-1 can lead to increased activation of the IGF-1 receptor (IGF-1R) and downstream signaling pathways. This may contribute to enhanced cellular proliferation, survival, and potentially hypertrophic responses in the context of the myocardium.

On the other hand, uncontrolled IGF-1 signaling, resulting from destabilized IGFBP-1, can potentially promote cardiomyocyte hypertrophy. This hypertrophic response serves as an adaptive mechanism to preserve cardiac function following a myocardial infarction (Heinen et al., 2019). This process enhances contractility and facilitates tissue remodeling to improve overall heart function in the aftermath of myocardial infarction (Macvanin, M., et al., 2023).

### PCSK9

PCSK9 is primarily known for its role in cholesterol homeostasis (Coudert E et al., 2023). It regulates the number of low-density lipoprotein receptors (LDLRs) on the surface of liver cells. LDLRs play a crucial role in removing LDL (low-density lipoprotein) from the bloodstream. PCSK9 acts by binding to LDLRs and promoting their degradation, reducing the liver’s ability to remove LDL from the blood (Lagace T. A., 2014).

During our computational analysis, we pinpointed a missense single nucleotide polymorphism (missense SNP) characterized by an amino acid change at position R93C. Notably, this mutation was associated with a decrease in the protein’s stability. This instability might be attributed to the mutation’s location within the peptidase S8 propeptide region, which acts as a molecular chaperone for protein folding and activation following propeptide cleavage (Paysan-Lafosse T., et al., 2022).

Interestingly, this particular missense SNP in PCSK9 does not appear to elevate the risk of myocardial infarction but rather shows potential for reducing such risk. This risk reduction is attributed to the mutation’s destabilizing effect on PCSK9, which can impede its interactions with key partner proteins like LDLR and VLDLR, consequently hampering the process of LDL removal from the bloodstream.

On the other hand, PCSK9 also interacts with another important protein, ANXA2, which has been identified as both an endogenous binding partner and a functional inhibitor of PCSK9 (Seidah NG., et al., 2012). Given that reduced PCSK9 stability may compromise its ability to interact with and bind to ANXA2, it implies a broader impact on the regulatory mechanisms involving PCSK9 and its associated proteins.

## 5. Conclusion

The comprehensive computational bioinformatics research, aimed at uncovering missense SNPs linked to myocardial infarction, has provided valuable insights into the genetic landscape influencing cardiac health. The investigation into ALDH2 illuminated a destabilizing missense SNP (5504K), prompting concerns about its impact on aldehyde detoxification and the cardioprotective role of this vital enzyme. APOE analysis revealed a missense SNP (R176C) associated with reduced stability, potentially disrupting lipid transport and metabolism, thus implicating APOE in conditions like atherosclerosis and myocardial infarction. IGFBP-1 analysis identified an E218V missense SNP with destabilizing potential, hinting at uncontrolled IGF-1 signaling and its role in cardiomyocyte hypertrophy post-myocardial infarction. Conversely, PCSK9 analysis uncovered an R93C missense SNP, indicating decreased stability without a direct link to contributing to myocardial infarction risk.

It is crucial to note that these findings are based on computational analyses, and their translational impact requires experimental validations to address inherent limitations and ensure clinical relevance. Nevertheless, this work lays a foundation for future research and potential therapeutic interventions in myocardial infarction prevention and management.

## Acknowledgments

This work is supported by the Summer Undergraduate Research Apprenticeship (SURA) under Carnegie Mellon University Qatar, Doha, Qatar.

## Author Contributions

F.T. performed in-silico analysis, utilized computational tools, prepared the figures, and reviewed the manuscript. S.R. guided with the project design and approved the manuscript.

## References

Badr EA, Mostafa RG, Awad SM, Marwan H, Abd El-Bary HM, Shehab HE, Ghanem SE. A pilot study on the relation between irisin single-nucleotide polymorphism and risk of myocardial infarction. Biochem Biophys Rep. 2020 Feb 21;22:100742. doi: 10.1016/j.bbrep.2020.100742. PMID: 32123756; PMCID: PMC7038008.

Bae, J. H., Song, D. K., & Im, S. S. (2013). Regulation of IGFBP-1 in Metabolic Diseases. Journal of lifestyle medicine,3(2), 73–79.

Budas, G. R., Disatnik, M. H., & Mochly-Rosen, D. (2009). Aldehyde dehydrogenase 2 in cardiac protection: a new therapeutic target?. Trends in cardiovascular medicine,19(5), 158–164. 10.1016/j.tcm.2009.09.003

Chen QF, Wang W, Huang Z, Huang DL, Li T, Wang F, Li J. Correlation of rs1122608 SNP with acute myocardial infarction susceptibility and clinical characteristics in a Chinese Han population: A casecontrol study. Anatol J Cardiol. 2018 Apr;19(4):249–258. doi: 10.14744/AnatolJCardiol.2018.35002. PMID: 29615549; PMCID: PMC5998852.

Coudert E, Gehant S, de Castro E, Pozzato M, Baratin D, Neto T, Sigrist C J A, Redaschi N, Bridge A, UniProt Consortium. Annotation of biologically relevant ligands in UniProtKB using ChEBI Bioinformatics39:btac793(2023)

Dai X, Wiernek S, Evans JP, Runge MS. Genetics of coronary artery disease and myocardial infarction. World J Cardiol. 2016 Jan 26;8(1):1–23. doi: 10.4330/wjc.v8.i1.1. PMID: 26839654; PMCID: PMC4728103.

Heinen, A., Nederlof, R., Panjwani, P., Spychala, A., Tschaidse, T., Reffelt, H., Boy, J., Raupach, A., Gödecke, S., Petzsch, P., Köhrer, K., Grandoch, M., Petz, A., Fischer, J. W., Alter, C., Vasilevska, J., Lang, P., & Gödecke, A. (2019). IGF1 Treatment Improves Cardiac Remodeling after Infarction by Targeting Myeloid Cells. Molecular therapy : the journal of the American Society of Gene Therapy,27(1), 46–58. 10.1016/j.ymthe.2018.10.020

Hopkins PN, Brinton EA, Nanjee MN. Hyperlipoproteinemia type 3: the forgotten phenotype. Curr Atheroscler Rep. 2014 Sep;16(9):440. doi: 10.1007/s11883-014-0440-2. PMID: 25079293.

Kim SJ. Global Awareness of Myocardial Infarction Symptoms in General Population. Korean Circ J. 2021 Dec;51(12):997–1000. doi: 10.4070/kcj.2021.0320. PMID: 34854579; PMCID: PMC8636759.

Lagace T. A. (2014). PCSK9 and LDLR degradation: regulatory mechanisms in circulation and in cells. Current opinion in lipidology,25(5), 387–393. 10.1097/MOL.0000000000000114

Li J, Qin R, Wang W, Huang Z, Huang DL, Li T, Wang F, Zeng XT, Sun ZY, Liu XF, Huang F, Guo T. Relationship between SNP rs1764391 and Susceptibility, Risk Factors, Gene-environment Interactions of Acute Myocardial Infarction in Guangxi Han Chinese Population. Curr Pharm Biotechnol. 2020;21(1):79–88. doi: 10.2174/1389201019666191003150015. PMID: 31580250.

Macvanin, M., Gluvic, Z., Radovanovic, J., Essack, M., Gao, X., & Isenovic, E. R. (2023). New insights on the cardiovascular effects of IGF-1. Frontiers in endocrinology, 14, 1142644. 10.3389/fendo.2023.1142644

Pan Q, Hui D, Hu C. A Variant of IL1B Is Associated with the Risk and Blood Lipid Levels of Myocardial Infarction in Eastern Chinese Individuals. Immunol Invest. 2022 Jul;51(5):1162–1169. doi: 10.1080/08820139.2021.1914081. Epub 2021 May 3. PMID: 33941028.

Paysan-Lafosse T, Blum M, Chuguransky S, Grego T, Pinto BL, Salazar GA, Bileschi ML, Bork P, Bridge A, Colwell L, Gough J, Haft DH, Letunić I, Marchler-Bauer A, Mi H, Natale DA, Orengo CA, Pandurangan AP, Rivoire C, Sigrist CJA, Sillitoe I, Thanki N, Thomas PD, Tosatto SCE, Wu CH, Bateman A. InterPro in 2022. Nucleic Acids Research, Nov 2022, (doi: 10.1093/nar/gkac993)

Saleh M, Ambrose JA. Understanding myocardial infarction. F1000Res. 2018 Sep 3;7:F1000 Faculty Rev-1378. doi: 10.12688/f1000research.15096.1. PMID: 30228871; PMCID: PMC6124376.

Seidah NG, Poirier S, Denis M, Parker R, Miao B, Mapelli C, Prat A, Wassef H, Davignon J, Hajjar KA, Mayer G. Annexin A2 is a natural extrahepatic inhibitor of the PCSK9-induced LDL receptor degradation. PLoS One. 2012;7(7):e41865. doi: 10.1371/journal.pone.0041865. Epub 2012 Jul 27. PMID: 22848640; PMCID: PMC3407131.

Semaev S, Shakhtshneider E, Orlov P, Ivanoshchuk D, Malyutina S, Gafarov V, Ragino Y, Voevoda M. Association of RS708272 (CETP Gene Variant) with Lipid Profile Parameters and the Risk of Myocardial Infarction in the White Population of Western Siberia. Biomolecules. 2019 Nov 14;9(11):739. doi: 10.3390/biom9110739. PMID: 31739638; PMCID: PMC6921014.

Škrlec I, Milić J, Steiner R. The Impact of the Circadian Genes CLOCK and ARNTL on Myocardial Infarction. J Clin Med. 2020 Feb 10;9(2):484. doi: 10.3390/jcm9020484. PMID: 32050674; PMCID: PMC7074039.

Szklarczyk D, Franceschini A, Wyder S, Forslund K, Heller D, Huerta-Cepas J, Simonovic M, Roth A, Santos A, Tsafou KP, Kuhn M, Bork P, Jensen LJ, von Mering C. STRING v10: protein-protein interaction networks, integrated over the tree of life. Nucleic Acids Res. 2015 Jan;43(Database issue):D447–52. doi: 10.1093/nar/gku1003. Epub 2014 Oct 28. PMID: 25352553; PMCID: PMC4383874.

U.S. National Library of Medicine. (n.d.). IGFBP1 insulin like growth factor binding protein 1 [homo sapiens (human)] - gene - NCBI. National Center for Biotechnology Information. https://www.ncbi.nlm.nih.gov/gene/3484

